# Detection and characterization of a novel copper-dependent intermediate in a lytic polysaccharide monooxygenase

**DOI:** 10.1101/610865

**Authors:** Raushan K. Singh, Bart v. Oort, Benedikt Möllers, David A. Russo, Ranjitha Singh, Høgni Weihe, Manish K. Tiwari, Roberta Croce, Paul E. Jensen, Claus Felby, Morten J. Bjerrum

## Abstract

Lytic polysaccharide monooxygenases (LPMOs) are copper-containing enzymes capable of oxidizing crystalline cellulose and the enzyme has large practical application in the process of refining biomass. The LPMO catalytic mechanism still remains debated despite several proposed reaction mechanisms. Here, we report a long-lived intermediate (*t*_½_ = 6 – 8 minutes) observed in an LPMO from *Thermoascus aurantiacus* (TaLPMO9A). The intermediate with a strong absorption around 420 nm is formed when reduced LPMO-Cu(I) reacts with H_2_O_2_. UV-vis absorption spectroscopy, electron paramagnetic resonance (EPR), and stopped-flow spectroscopy indicate that the observed long-lived intermediate involves the copper center and a nearby tyrosine (Tyr175). We propose that the reaction with H_2_O_2_ first forms a highly reactive short-lived Cu(III)-intermediate which is subsequently transformed into the observed long-lived copper-dependent intermediate. Since sub-equimolar amount of H_2_O_2_ to LPMO boosts oxidation of phosphoric acid swollen cellulose (PASC) suggests that the long-lived copper-dependent intermediate is part of the catalytic mechanism for LPMOs. The proposed mechanism offers new perspectives in the oxidative reaction mechanism of copper enzymes and hence for the biomass oxidation and the reactivity of copper in biological systems.

## INTRODUCTION

The cleavage of carbon-hydrogen bonds is central to the biological breakdown of organic matter into carbon dioxide and water. With the discovery of lytic polysaccharide monooxygenases (LPMOs), it was shown for the first time, how an enzyme with a single copper in the active site can activate dioxygen to break carbon-hydrogen bonds and degrade the most abundant and recalcitrant polysaccharides, such as cellulose and chitin ^1^. Since their discovery, LPMOs have been found in fungi, bacteria, virus, invertebrates, and algae, highlighting their biological importance ^2^. The diversity of identified LPMOs has increased with currently six different families classified in the CAZY database as auxiliary activity (AA) AA9, AA10, AA11, AA13, AA14 and AA15 families ^3,4^. However, despite the importance of LPMOs and the extensive research in their structure and enzymology, the actual catalytic mechanism is yet to be resolved.

LPMOs have a solvent-exposed mononuclear type-2 copper active site, with a t-shaped coordination sphere named the histidine–brace, where two histidine residues provide three nitrogen ligands, two from N-His and one from the terminal amine ^5^. Another characteristic of LPMOs is that the copper also coordinates to an adjacent aromatic amino acid; for AA9, AA11, AA13, AA14, and AA15 LPMOs the amino acid is a tyrosine, whereas for AA10 LPMOs it is mostly a phenylalanine with few exceptions ^6^. The overall reaction of LPMOs has been proposed to be the abstraction of a hydrogen atom from the glycoside C-H bond followed by insertion of an oxygen atom in the carbohydrate chain and thereby cleaving the glycosidic bond ^7^. Specifically, for glucans, the oxidation of the pyranose ring can take place at either the C1 position producing aldonic acids or at the C4 position producing 4-ketoaldose (gemdiols) ^8^. The catalytic cycle is believed to be initiated by the reduction of LPMO-Cu(II) to LPMO-Cu(I), by an external electron donor, followed by binding of dioxygen to LPMO-Cu(I) ^9^. The nature of the external electron donor varies widely and four LPMO electron donor systems have been reported: fungal metabolites, lignin/phenols, GMC oxidoreductases and photoexcited pigments ^10–15^. The function of LPMOs has been subject to great controversy ^15–18^. In the currently established model LPMOs are seen as monooxygenases. However, this view is challenged by new data indicating that both O_2_ and H_2_O_2_ can function as co-substrates ^15,17,18^. This is further complicated by the fact that LPMOs can generate H_2_O_2_ when the enzyme is not bound to the substrate, a pathway referred to as the “futile cycle” ^15^. A copper-bound oxygen intermediate has been shown by neutron scattering ^7,19,20^, but the nature of the oxygen intermediate remains unknown. Kjaergaard et al. (2014) using DFT modelling suggested the presence of a [CuO_2_]^+^ superoxide species. Other theoretical works have shown that the superoxide in [CuO_2_]^+^ is not sufficiently reactive to abstract a hydrogen from the glycoside C–H bond ^19,21^ and have suggested that copper-oxyl, [CuO]^+^ copper-hydroxy, [CuOH]^2+^ or copper-hydroperoxide [CuOOH] species ^7,22–26^ may be generated by reductions and protonations of e.g. the superoxide in [CuO_2_]^+^. Walton and Davis analyzed a wide range of possible catalytic cycles with different oxygen intermediates and divided them into two classes based on the LPMO being free or bound to a substrate ^22,27^. They also indicated that some of the suggested pathways could involve Cu(III) as a reactive species.

The possible formation of Cu(III) in mono-nuclear copper sites is supported by earlier studies. The oxidation of copper complexes with tri- and tetrapeptides has been reported by Margerum and co-workers ^28–30^, showing that it was possible to form Cu(III) complexes with a lifetime of several minutes. Cu(III) complexes generated by mixing the copper-coordinated tetrapeptide (Cu(II)Gly2HisGly) with ascorbate, and H_2_O_2_ decomposed to relatively stable alkene peptides ^30^. The oxidation could also be performed electrochemically or by strong oxidants (e.g. [Ir(IV)Cl_6_]^2−^) ^30^. The formed copper(III)-tetrapeptide showed UV-vis absorption around 400 nm with a molar absorbance of 1.7·10^3^ M^−1^cm^−1 30^. Comparing the ligand environment of the Cu(II) site in the tetrapeptides with that of the Cu(II) site in LPMOs shows that the Cu(II) sites in LPMOs have large structural similarity with these simple Cu(II) peptide complexes. A major difference between the structure of LPMOs ^5^ and the copper(II)-tetrapeptide complexes is that the Cu(II) in LPMOs is interacting with the nearby aromatic amino acid (tyrosine (Tyr175) in TaLPMO9A). Interestingly, tyrosine radical is reported to have molar absorbance of 2.5·10^3^ – 5.0·10^3^ at 420 nm ^31^. It is also worth mentioning that ascorbate has been used as redox cofactor and H_2_O_2_ as co-substrate for LPMO-catalyzed oxidation ^5,15,32^.

It was therefore investigated if a possible Cu(III) and/or tyrosine radical species were formed when LPMOs was treated with various combinations of ascorbate and H_2_O_2_. A new LPMO intermediate with absorption around 420 nm involving the active site copper and a nearby tyrosine was observed and its reactivity, formation and decay were studied using spectroscopic and chromatographic tools.

## RESULTS

### Oxidation of TaLPMO9A-Cu(II) by ascorbate and H_2_O_2_

To test if copper in LPMOs could be oxidized to Cu(III) as seen for copper coordinated to tri- and tetrapeptides ^30^, we oxidized the LPMO by ascorbate and H_2_O_2_. For this study, we chose a well-characterized AA9 LPMO from *Thermoascus aurantiacus* (TaLPMO9A) also known as TaGH61A/TaAA9A ^5^. Highly pure and fully Cu(II) loaded *holo*TaLPMO9A (Cu/LPMO molar ratio is 1.07 ± 0.06) was obtained as outlined in Materials and Methods (Fig. S1). Impurities from the protein including media impurities were removed and the enzyme was highly pure as judged from SDS-PAGE (see Fig. S1B). UV-vis spectra (Fig. 1) shows the changes occurring when purified *holo*TaLPMO9A is mixed with ascorbate followed by the addition of sub-equimolar amounts of H_2_O_2_ (7:1 protein: H_2_O_2_ ratio). The UV-vis spectra shows an absorption peak appearing at 420 nm after addition of ascorbate and H_2_O_2_ (Fig. 1A); this peak decreases with time, having nearly disappeared after 45 min. (Fig. 1A and 1B). In this experiment, the 420 nm peak decay is close to a first-order decay with a half-life of approximately 6 – 8 minutes (Fig. 1B). As shown in Fig. S2, a decrease in Cu(II) absorbance around 640 nm and in the CD signal around 720 nm after addition of ascorbate confirmed that the LPMO-Cu(II) was partially reduced to LPMO-Cu(I) before the addition of H_2_O_2_. The 420 nm peak is not observed when the *apo*LPMO is treated with ascorbate and H_2_O_2_ (Fig. 2). Furthermore, no peak is observed at 420 nm when *holo*TaLPMO9A is oxidized with H_2_O_2_ in the absence of ascorbate or when H_2_O_2_ is added to *holo*TaLPMO9A *before* addition of ascorbate. This suggests that the presence of copper is essential and that LPMO-Cu(II) *cannot* form the product absorbing at 420 nm by reaction with H_2_O_2_ alone, and that the reduction of LPMO-Cu(II) to LPMO-Cu(I) is essential for the formation of the 420 nm peak. Margerum and co-workers have also reported the lack of oxidation of Cu(II)-(H-2Gly2HisGly)**-**to Cu(III) with H_2_O_2_ alone ^30^.

**Fig. 1.**
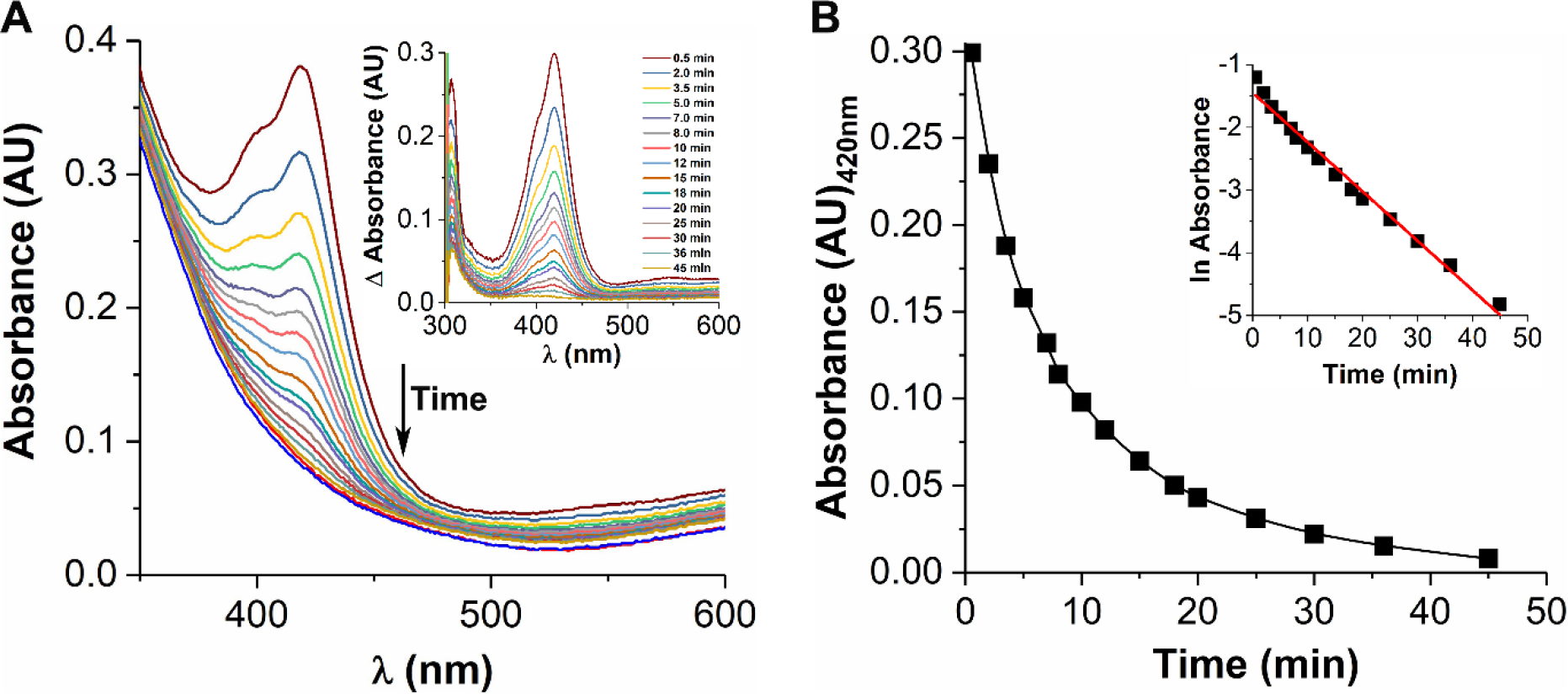
UV-vis spectral change when TaLPMO9A is mixed with ascorbate and H_2_O_2_. (**A**) UV-vis spectra of reduced TaLPMO9A after addition of sub-equimolar concentrations of H_2_O_2_. Red and blue lines represent the *holo*TaLPMO9A and reduced *holo*TaLPMO9A spectra, respectively. After addition of H_2_O_2_ to reduced *holo*TaLPMO9A peak at 420 nm appears which decays with time. The inset shows the peak at 420 nm at a different time intervals after subtracting the reduced *holo*TaLPMO9A spectra from reduced *holo*TaLPMO9A + H_2_O_2_ spectra. (**B**) Decay of the absorption peak at 420 nm over time. The inset shows the linear plot of ln absorbance over time. Experimental conditions: TaLPMO9A 1.39 mM, ascorbate 2 mM, H_2_O_2_ 0.2 mM in 20 mM phosphate buffer at pH 6.6 and room temperature.

**Fig. 2.**
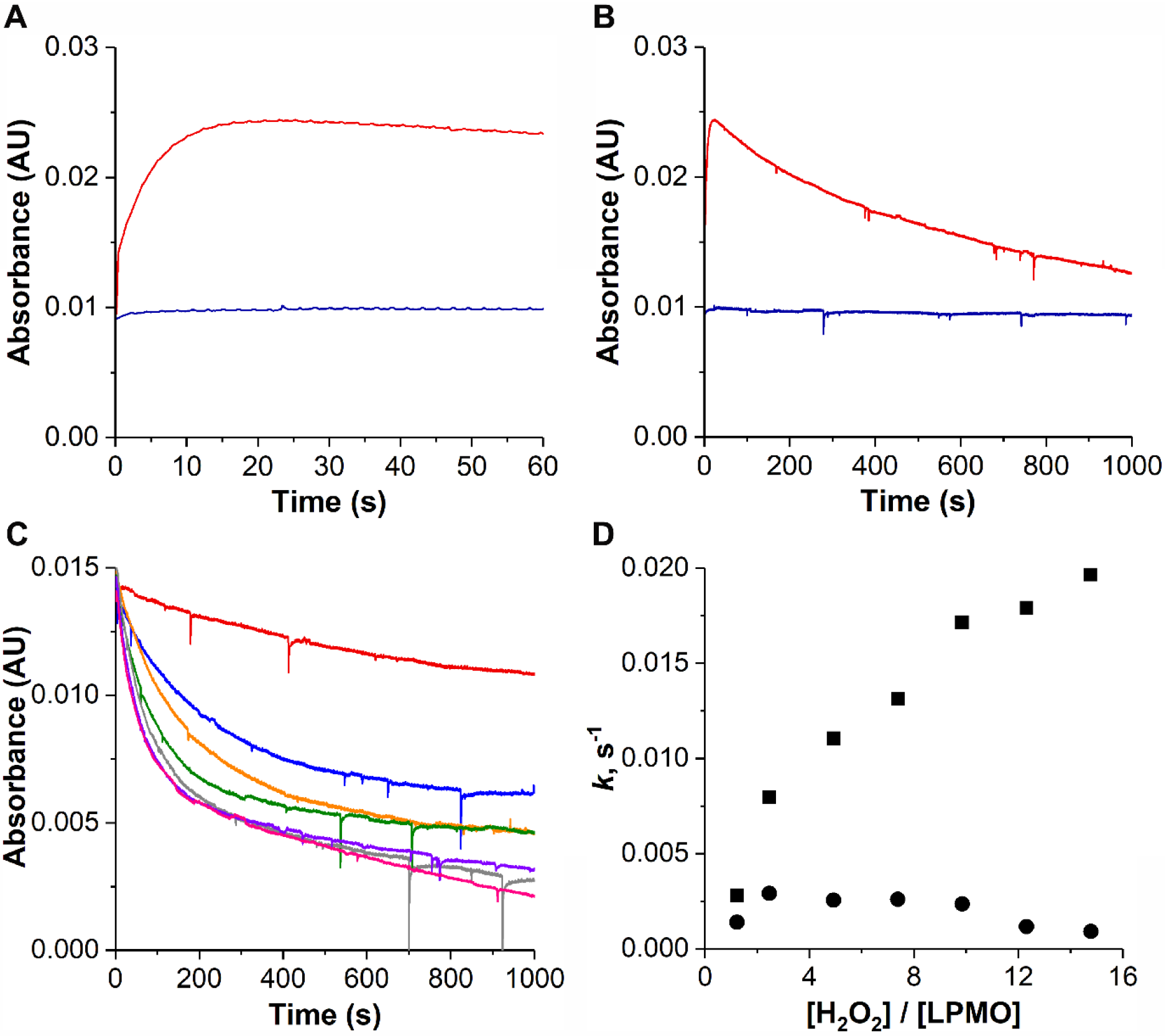
Single wavelength absorption transients at 420 nm for the reaction of reduced TaLPMO9A with H_2_O_2_. Transients absorbance observed for 60 s (**A**) and 1000 s after the addition of H_2_O_2_ (**B**). Red and blue lines represent the *holo*TaLPMO9A and *apo*TaLPMO9A, respectively. Experimental conditions: *apo*TaLPMO9A or *holo*TaLPMO9A 0.2 mM, ascorbate 1 mM, H_2_O_2_ 0.1 mM in 20 mM phosphate buffer at pH 6.6 and 20 °C. (**C**) Transients absorbance during the reaction of reduced TaLPMO9A with varying concentrations of H_2_O_2_ observed for 1000 s. Red, blue, orange, green, violet, grey and pink lines represent 0.25, 0.5, 1.0, 1.5, 2.0, 2.5 and 3.0 mM H_2_O_2_, respectively. Rate constants were determined by fitting the transients to equation 2 (Materials and methods) and are given in the supplementary information. Fitting was performed for the time period 5 to 1000 s. Absorbance was normalized to same peak intensity and plotted against time (**D**). Fitted rate constants as a function of varying H_2_O_2_ concentrations for reduced TaLPMO9A. Rates, *k*_1_ (fast) and *k*_2_ (slow) represented as solid square and solid circle, respectively. Only data collected after 5s were analyzed and fitted. Experimental conditions: *holo*TaLPMO9A 0.2 mM, ascorbate 1 mM in 20 mM phosphate buffer at pH 6.6 and 20 °C.

### Formation and decay kinetics of intermediate

The formation and decay of the intermediate were investigated by stopped-flow kinetics. By addition of 0.1 mM H_2_O_2_ (2:1 protein: H_2_O_2_ ratio) the intermediate is formed in less than 10 s and then gradually decays with a lifetime of around 6 – 8 minutes (Fig. 2A and 2B). For *apo*TaLPMO9A (i.e. copper free), no intermediate is observed (Fig. 2A and 2B). Subsequent stopped-flow experiments at varying concentrations of H_2_O_2_ (Fig. 2C) show that increasing amounts of H_2_O_2_ reduces the formation and lifetime of the intermediate. A simple bi-exponential fit to the data indicates that the decay reaction contains both a first- and a second-order component (Fig. 2D, Table S1). This suggests that the decay proceeds both as a “second-order” reaction with H_2_O_2_ (*k*_1_ increases with H_2_O_2_ concentration) and as a “first-order” decay (*k*_2_). Furthermore, pH variation (5.9 to 7.6) had only small effect on the formation and decay of the intermediate (Fig. S3).

### Paramagnetic behaviour of intermediate

The intermediate, formed at sub-equimolar amounts of H_2_O_2_ had a lifetime of around 6 – 8 minutes. It is therefore possible to investigate the nature of the intermediate using EPR spectroscopy. The EPR spectrum of the *holo*TaLPMO9A/ascorbate/H_2_O_2_ system was obtained at 77 K (Fig. S4). The EPR spectra of partly reduced TaLPMO9A and partly reduced TaLPMO9A with addition of 0.5 equivalents H_2_O_2_ were essentially identical to those of TaLPMO9A (Fig. S4). The parameters for the EPR spectra at 77K are given in Table S2. The observed EPR spectrum is characteristic of a type 2 copper site with *g*_z_ (*g*_║_) =2.27 and *g*_xy_ (*g*┴) = 2.06 and *A*_z_ (*A*_║_) = 492 MHz. Furthermore, addition of ascorbate reduced the signal without changing the obtained parameters (Table S2).

Next, to mimic the conditions in which we observed the intermediate at 420 nm in the UV-vis experiment, the EPR spectra were measured at room temperature. The spectrum of *holo*TaLPMO9A in solution, in the presence of both ascorbate and ascorbate/H_2_O_2_ is shown in Fig. 3A and the fit parameters are given in Table 1. The parameters *g*_z_ = 2.27 and *g*_xy_= 2.06 and *A*_z_ = 459/MHz are close to those obtained at 77 K. Furthermore, in correlation with the reduction of TaLPMO9A-Cu(II) to TaLPMO9A-Cu(I) addition of ascorbate reduced the signal to about half intensity without changing the obtained parameters as seen from Cu(conc.) in Table 1. The EPR spectrum after addition of both ascorbate and H_2_O_2_ has an increased signal with some broadening. The difference in EPR signal in Fig. 3B is obtained by subtracting the reduced TaLPMO9A spectra (blue) from the spectrum of the ascorbate/H_2_O_2_ treated TaLPMO9A (green). The parameters for this difference spectrum are *g*_z_ = 2.17 and *g*_xy_ = 2.07.

**Table 1.** EPR spectral parameters for TaLPMO9A at room temperature. I, and II denote TaLPMO9A, and reduced TaLPMO9A, respectively. IV denotes the EPR parameters of the **s**pectra after subtracting the reduced TaLPMO9A spectra from H_2_O_2_ treated reduced TaLPMO9A spectra.

| Parameters | I | II | IV |
| --- | --- | --- | --- |
| Cu (A<sub>z</sub>) (MHz) | 459(1) | 459(2) | - |
| Cu (g<sub>xy</sub>) | 2.0637(1) | 2.0638(2) | - |
| Cu (g<sub>z</sub>) | 2.2670(3) | 2.2658(4) | - |
| Cu (conc.) | 852(17) | 409(15) | - |
| S=1 (g<sub>xy</sub>) | - | - | 2.067(2) |
| S=1 (g<sub>z</sub>) | - | - | 2.17(1) |
| S=1 (conc.) | - | - | 636(37) |

**Fig. 3.**
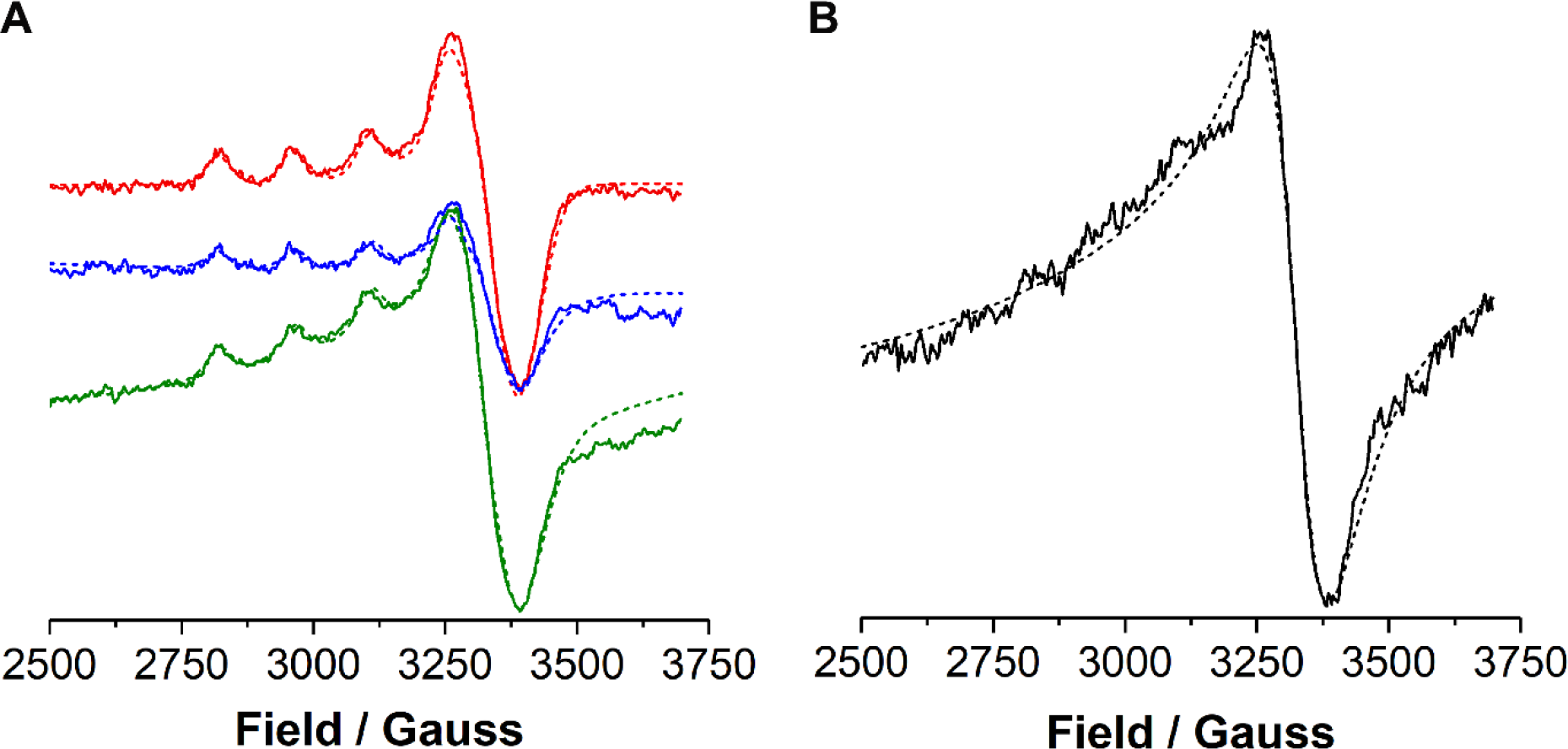
EPR spectra of *holo*TaLPMO9A. (**A**) X-band EPR spectra of TaLPMO9A (red), reduced TaLPMO9A (blue) and H_2_O_2_ treated reduced TaLPMO9A (green) at room temperature in 20 mM phosphate buffer, pH 6.6. The sample was put in a capillary and then carefully placed inside the quartz EPR tube. The spectrum recorded for the buffer in the capillary was used as the reference to remove the background signal. (**B**) Spectra after subtracting the TaLPMO9A spectra from H_2_O_2_ treated reduced TaLPMO9A spectra. Solid lines indicate experimental spectra recorded of solutions at room temperature. The dashed lines indicate computed spectra obtained with parameters reported in Table 1. Experimental conditions: 3.6 mM TaLPMO9A, 4 mM ascorbate, and 2 mM H_2_O_2_ at pH 6.6 in 20 mM phosphate buffer.

### Influence of the intermediate formation on LPMO activity

To elucidate the potential role of the observed intermediate in the catalytic cycle of the LPMO, we used a substrate conversion assay with PASC as substrate. The released oligomeric products were detected and quantified by high-performance anion exchange chromatography (HPAEC) (Fig. 4). At 0.1 mM H_2_O_2_, the release of non-oxidized oligosaccharides derived from PASC oxidation was enhanced whereas the release of C4-oxidized products only showed minor differences relative to a sample without addition of exogenous H_2_O_2_ (Fig. 4A). The correlation to the mechanism when only O_2_ is present is further elaborated in the discussion. As previously observed from Stopped-flow experiments, the lifetime of the intermediate is approximately inversely correlated with the concentration of H_2_O_2_. We therefore, proceeded with experiments in the presence of 20 times higher H_2_O_2_ concentration (Fig. 4B). Interestingly, only a negligible release of C4 oxidation products was detected which is in accordance with the level of observed intermediate.

**Fig. 4.**
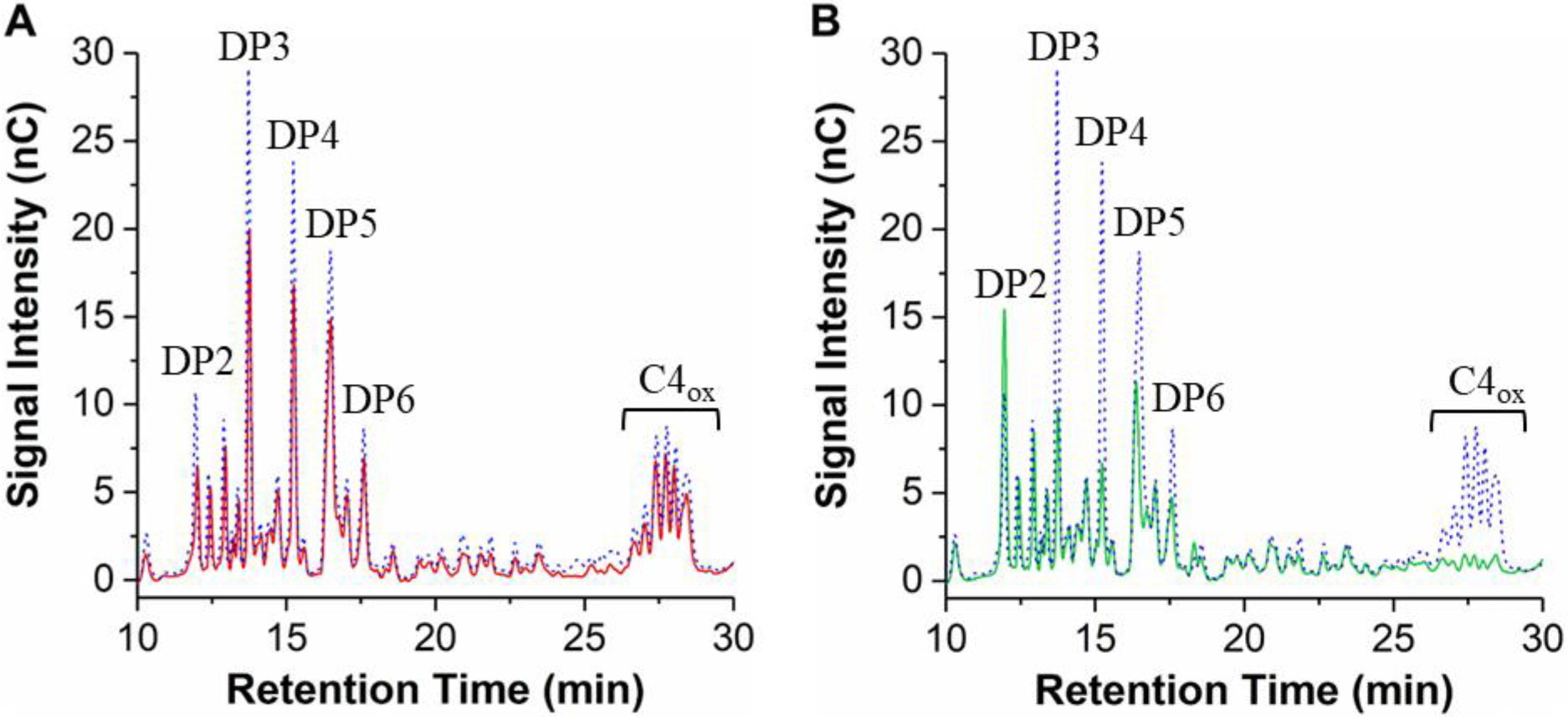
TaLPMO9A catalyzed oxidation of PASC. (**A**) HPAEC chromatogram showing the TaLPMO9A catalyzed oxidation of PASC with supplementation of 0.1 mM H_2_O_2_ (blue) and without H_2_O_2_ (red). (**B**) TaLPMO9A catalyzed oxidation of PASC was supplemented with 0.1 mM H_2_O_2_ (blue) and 2.0 mM H_2_O_2_ (green). Experimental conditions: TaLPMO9A (0.1 mM), PASC (0.4 %), ascorbate (2 mM), in phosphate buffer (20 mM, pH 6.0) at 25 °C and 800 rpm for 30 min. The peaks were assigned according to the elution profile of commercially available non-oxidized cello-oligosaccharides. DP2, cellobiose; DP3, cellotriose; DP4, cellotetraose; DP5, cellopentaose; DP6, cellohexaose. The C4-oxidized glucose (4-keto aldose) peaks appear after a retention time of 25 min.

### LPMO oxidation and dimerization

Finally, the post-assay samples were analyzed by SDS-PAGE, which revealed dimer formation of the TaLPMO9A (Fig. 5A). The dimerization can only be observed with the complete *holo*TaLPMO9A/ascorbate/H_2_O_2_ system and no dimers were formed in reaction mixtures where only LPMO and ascorbate were present, or in the absence of Cu (*apo*LPMO) (Fig. 5A, Fig. S5A, Fig. S5B and Fig. 6B). The LPMO used in this study has 12 tyrosine residues and their positions are shown in the X-ray crystal structure in Fig. S5C ^5^. It is conceivable that the observed dimerization could be due to intermolecular dityrosine formation. To confirm this hypothesis, we followed the fluorescence at 405 nm, the emission wavelength expected for dityrosine ^33^, over a period of 24 hours (Fig. 5B). The observed increase in fluorescence over time strongly indicates that the dimer formation is, in fact, due to dityrosine cross-linking between LPMOs in solution. Interestingly, this dimer formation is only seen under conditions where the intermediate is observed and occurs over a period of 24 hours which is much slower than the decay of the intermediate (Fig. 1B). Based on the observation from SDS-PAGE that the formation of the dimers is between 10 to 15 % of the whole LPMO sample it is estimated that the molar absorbance of the LPMO intermediate at 420 nm is between 1400 to 2200 M^−1^cm^−1^ (using data from Fig. 1A and 5A), assuming that two intermediate will result in one dimer. Which is within the range of other reported values on similar systems ^31^.

**Fig. 5.**
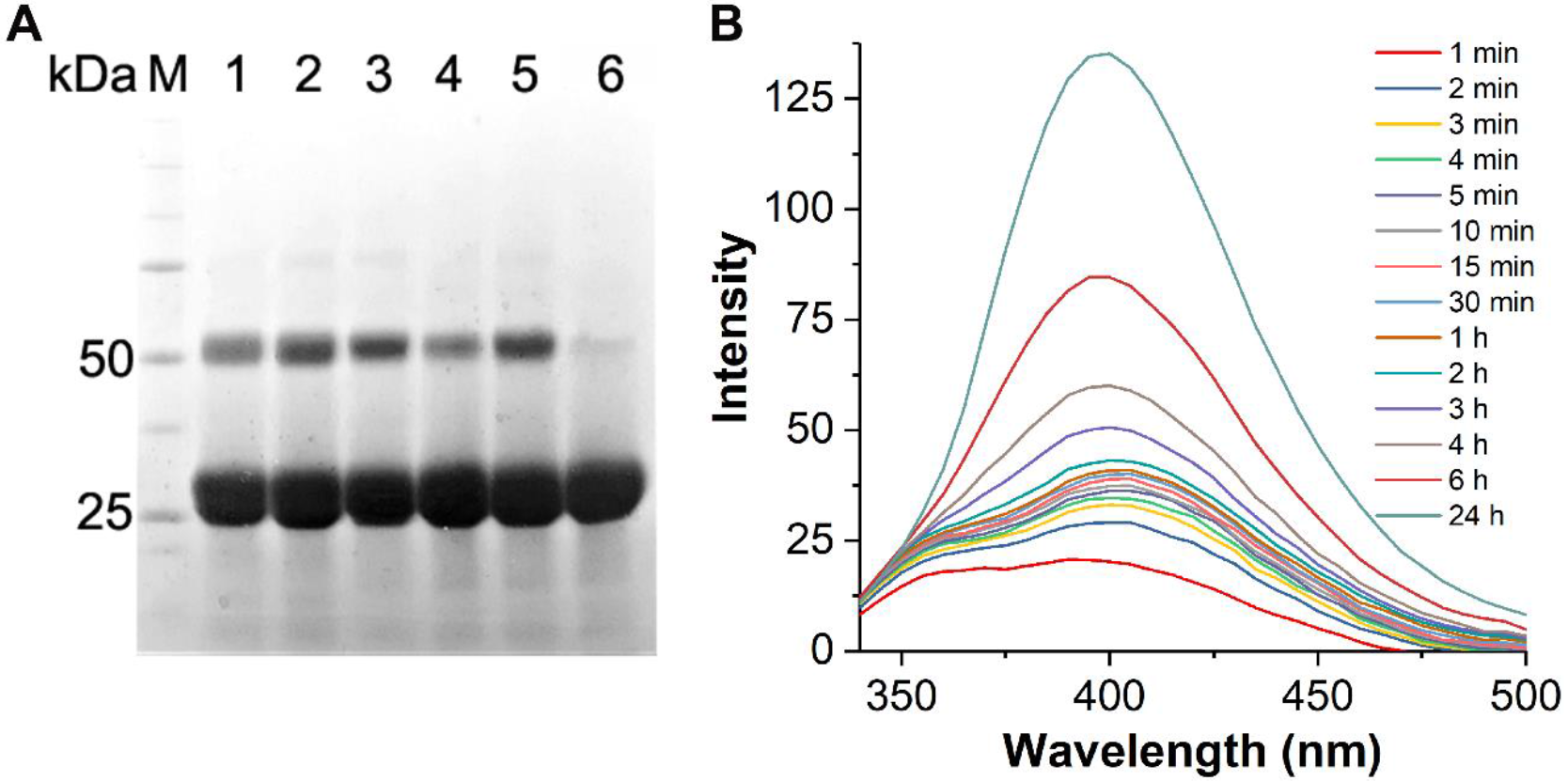
The appearance of TaLPMO9A dimers correlates with the formation of dityrosine. (**A**) SDS-PAGE analysis of the TaLPMO9A after initial intermediate formation. Lane 1: TaLPMO9A + 2 mM ascorbate + 2 mM H_2_O_2_, Lane 2: TaLPMO9A + 1 mM ascorbate + 1 mM H_2_O_2_, Lane 3: TaLPMO9A + 1 mM ascorbate + 0.5 mM H_2_O_2_, Lane 4: TaLPMO9A + 0.5 mM ascorbate + 0.2 mM H_2_O_2_, Lane 5: TaLPMO9A + 1 mM ascorbate + 0.2 mM H_2_O_2_, Lane 6: TaLPMO9A (1.39 mM). (**B**) Dityrosine formation followed by fluorescence spectroscopy with emission at 340-500 nm and excitation at 320 nm. The difference in fluorescence spectra is obtained by subtracting the reduced TaLPMO9A spectra from H_2_O_2_ treated reduced TaLPMO9A spectra. Assay was performed on Jasco FP-6300 spectrofluorometer. Experimental conditions: TaLPMO9A (1.05 mM) + ascorbate (2 mM) + H_2_O_2_ (0.5 mM) incubated in phosphate buffer (20mM, pH 6.6) at room temperature.

**Fig. 6.**
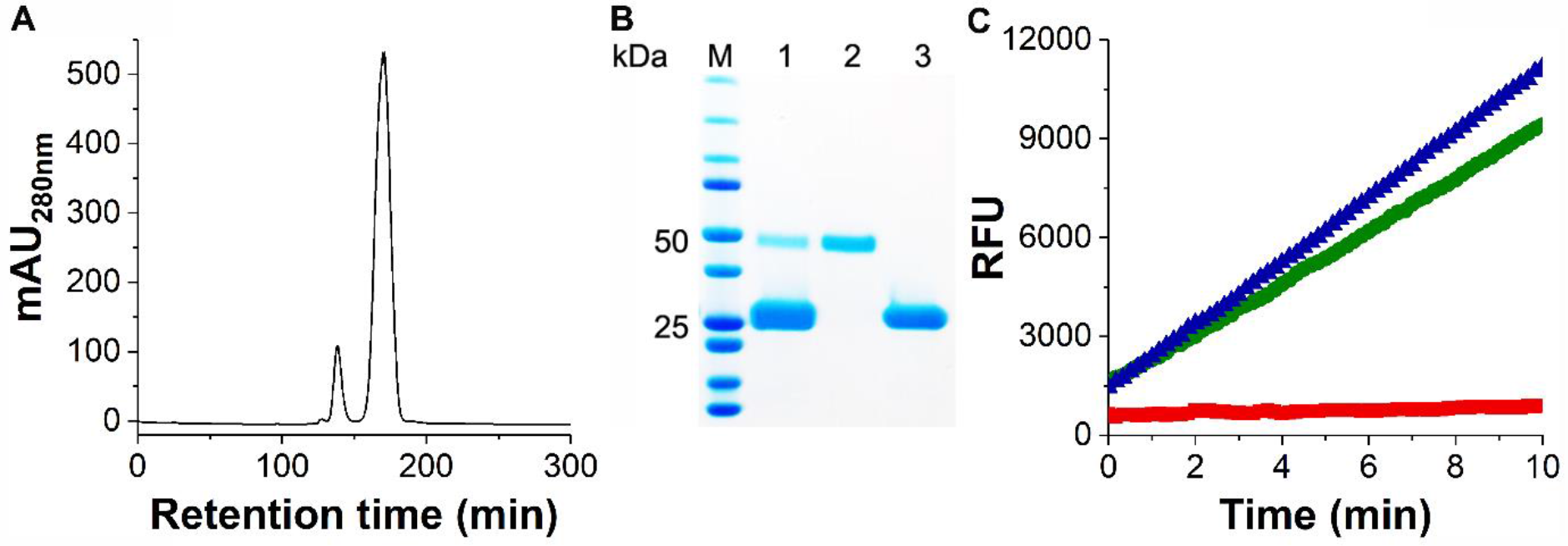
Purification of observed dityrosine dimers of TaLPMO9A and the activity as measured by Amplex red assay. (**A**) Chromatogram showing purification of oxidized TaLPMO9A using Hiload 26/60 Superdex 75 prep grade column. Oxidized TaLPMO9A was washed and concentrated before loading into the size exclusion column. Monomer and dimer peaks were well separated and eluted separately. (**B**) SDS-PAGE analysis of the purified TaLPMO9A after oxidation. Lane 1, oxidized TaLPMO9A, lane 2, purified dimer LPMO, lane 3, purified monomer LPMO. (**C**) Amplex red assay was performed using purified dimer and monomer LPMO. Green 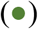 and blue 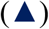 symbols represent activity for purified dimer and purified monomer LPMO, respectively. Red 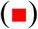 symbols represent reaction mixture without LPMO. The sample composition was: Ascorbate (0.08 mM), EDTA (0.1 mM), Amplex red (0.05 mM), HRP (20 U), purified-TaLPMO9A (2.0 μg/mL). The assay was performed on SpectraMax M2 multi-detection microplate reader using excitation and emission at 557 and 583 nm.

To probe if dimer formation had an effect on enzymatic activity, the LPMO dimers were purified by size exclusion chromatography (Fig. 6A and B). The two separate peaks observed by size exclusion chromatography and the two bands observed on SDS-PAGE clearly indicate that the oxidized TaLPMO9A exhibits intermolecular cross-linkage. Furthermore, the purified monomeric and dimeric fractions were tested for activity with an amplex red assay. The dimers maintained ~ 90% of the monomer activity (Fig. 6C). Furthermore, the peroxidase activity of TaLPMO9A was checked using 2,6-dimethoxyphenol (2,6-DMP) assay ^34^. The 2,6-DMP assay with TaLPMO9A showed that H_2_O_2_ could activate TaLPMO9A (Fig. S6). As shown in Fig. S6, the recorded spectra show the appearance of a peak right after the initiation of the reaction showing maxima at 469 nm (final reaction product coerulignone). This peak at 469 nm was less prominent (80-fold less OD_469nm_), when the reaction was performed in the absence of H_2_O_2_ (Fig. S6B and S6C). We also checked the peroxidase activity of oxidized TaLPMO9A (Fig. S8). Similarly, the peroxidase activity of oxidized TaLPMO9A (for TaLPMO9A oxidation, the reduced TaLPMO9A was treated with 0.5 and 10 eq of H_2_O_2_, and then purified before checking the peroxidase activity) retained 91 % and 78 % of its activity.

## DISCUSSION

The observed long-lived intermediate seen in the UV-Vis spectra of TaLPMO9A at 420 nm (Fig. 1), is formed by treating TaLPMO9A solution with ascorbate and H_2_O_2_. The presence of copper at the active site of TaLPMO9A is essential for the formation of the intermediate since no intermediate is formed when *apo*TaLPMO9A is used in the experiment (Fig. 2A and B). Treating *holo*TaLPMO9A with 1 mM ascorbate solution results in partial reduction of LPMO-Cu(II) to LPMO-Cu(I) as seen from the UV-vis and EPR measurements of TaLPMO9A with and without ascorbate (Fig. S2 and Fig. 3). Furthermore, the stopped-flow experiments (Fig.

2) showed that approximately 0.5 equivalents of H_2_O_2_ seem to give the highest observed level of the intermediate when using 1 mM ascorbate. Additionally, the *holo*TaLPMO9A upon mixing with H_2_O_2_ in the absence of ascorbate, does not show any intermediate formation. Combining the two observations, this is a strong indication that H_2_O_2_ only reacts with TaLPMO9A-Cu(I) during the formation of the intermediate. It is, therefore, possible that TaLPMO9A-Cu(I) with H_2_O_2_ forms LPMO-Cu(III). This conclusion is supported by Margerum and coworkers observation of Cu(III) formation when they treated tri- and tetrapeptides of Cu(II) with strong oxidants or ascorbate/H_2_O_2_ ^28–30^. It is important to note that the absorption around 420 nm from the LPMO intermediate disappears completely with the decay of the intermediate. Indicating that we do not see absorbance from stable oxidation products of TaLPMO9A like alkenes as described by Margerum and co-workers ^30^. However, we do observe the formation of dityrosine. This indicates that the decay mechanism of a Cu(III) species is different in the LPMOs compared to what happens when Cu(III) is formed in simple tri- and tetrapeptides. The active site in LPMO is somehow protected from oxidation by the possibly formed Cu(III).

The intermediate forms within seconds after mixing H_2_O_2_ with reduced TaLPMO9A as seen from the stopped-flow experiments (Fig. 2A). When no substrate and no surplus H_2_O_2_ is present, the intermediate is quite stable with a half-life of 6 – 8 min (Fig. 1). However, stopped-flow experiments with increasing amount of H_2_O_2_ clearly showed that the intermediate not only forms faster but decays even faster (Fig. 2C). The bi-exponential fit to the stopped-flow data indicated that the decay reaction contains both a first-order component as well as a second-order component proportional to the added H_2_O_2_ (Fig. 2D, Table S1) even if these data are not collected under strict pseudo-first order conditions. The average value (2.0·10^−3^ s^−1^ (*k*_2_)) for the first-order reaction obtained from stopped-flow experiment fits well with the value of 1.7·10^−3^ s^−1^ obtained from data in Fig. 1. Furthermore, the second-order decay is dominant with surplus H_2_O_2_ which explains why this intermediate has been so difficult to observe.

The chemical nature of the observed intermediate was investigated by X-band EPR spectroscopy (Fig. S4). It is clear that the 77 K EPR spectrum of TaLPMO9A is as expected when comparing with spectra already published in the literature ^5,35^. EPR measurements were also performed at room temperature to mimic conditions as close as possible to the stopped-flow experiments. Addition of ascorbate reduces the intensity of the EPR signal similar to what is expected when approximately half of the Cu(II) in the enzyme is reduced to Cu(I). Addition of 0.5 equivalents H_2_O_2_ to the partly reduced TaLPMO9A showed an increase in the EPR signal, and some line broadening (Fig. 3A). This could be indicative of a compound with an *S* = 1 signal growing up underneath the Cu(II) signal ^31^. Fitting of composite spectra with overlapping signals resulted, as expected, in poorly determined and highly correlated parameters. To quantify the concentration of the *S* = 1 species better, we subtracted the blue trace from the green trace resulting in the broad signal shown in Fig. 3B. Subtracting the two spectra gave an EPR signal (Fig. 3B) that was fitted with *g*_z_ = 2.17 and *g*_xy_= 2.07. This EPR signal is probably due to a ferromagnetically coupled triplet state arising in the TaLPMO9A from exchange coupling of two unpaired spins. Notice that the average value, 2.10(2), of the *g* factors for this species, is lower than that, 2.1311(4), of the Cu^2+^ signal. This is in agreement with the interpretation of this as originating from an *S* = 1 multiplet being a result of an interaction of Cu^2+^ with an organic radical. Canters and co-workers studied a radical intermediate in a small laccase (SLAC) formed in the presence of oxygen and ascorbate giving rise to an absorption band at 410 nm ^31^. The spin system was identified as a tyrosine radical ferromagnetically coupled to a Cu(II) giving rise to a triplet state. The active tyrosine was identified from the crystal structure of the SLAC as a Tyr108 radical coupled to the spin half of the T2 Cu in SLAC positioned approximately 5 Å away ^31,36,37^. Based on the similarity with the intermediate observed in SLAC, we propose that the observed intermediate of TaLPMO9A is due to the Cu-ion in the active site of the protein interacting with Tyr175 positioned approximately 2.79 Å away from it (Fig. S7) ^5,22,38^. Given the shorter distance compared to SLAC, it is conceivable to have favourable interaction between the active site copper and Tyr175. This suggests that the observed long-lived intermediate is actually a CuTyr-intermediate. A possible explanation for the formation of this CuTyr-intermediate could be that LPMO-Cu(I) is reacting with H_2_O_2_ forming a short-lived LPMO-Cu(III) (and 2 HO^−^). This LPMO-Cu(III) would likely coordinate with the proximate Tyr175 and form a Cu(III) combined with tyrosyl deprotonation. Such a structure is isoelectronic with a Cu(II) interaction with a tyrosyl radical and this resonance will stabilize the system and could explain the broad and weak EPR signal observed underneath the Cu(II) signal. An EPR signal is only expected from the Cu(II)–^•^OTyr structure since Cu(III) and a deprotonated tyrosyl in Cu(III) – ^⁃^OTyr do not have unpaired electrons. The reason why an EPR signal is not seen at 77 K could be that the equilibrium is shifted towards the Cu(III)–^⁃^OTyr structure and/or that the interaction changes in favor of an anti ferromagnetically coupling to Cu(II) at low temperature.

The kinetic data in Fig. 2C shows that the process for the disappearance of the CuTyr-intermediate can occur via two different pathways. The observed apparent first-order process is slow with a first-order half-life of 6 – 8 minutes. Treatment of reduced TaLPMO9A with sub-equimolar H_2_O_2_ reduce the enzyme activity slightly since the amplex red assay and 2,6-DMP assay showed more than 90% activity (Fig. 6 and Fig. S8A). The observed apparent second order process causing a decay of the intermediate in TaLPMO9A is dependent on the concentration of H_2_O_2_. This process seems to be comparatively more detrimental than the first order process since the activity of the TaLPMO9A after treatment with 10 eq of H_2_O_2_ reduced to 78 % (see Fig. S8B). The decay pathway of the CuTyr-intermediate is somehow correlated with the formation of dityrosines since dityrosin formation only is observed when the CuTyr-intermediate is formed. The dityrosine could be formed due to intra- or inter-molecular covalent linkage resulting in a TaLPMO9A dimer ^8,39,40^. Our observations indicates that H_2_O_2_ can indeed be a substrate for the LPMOs but only at low concentrations. As soon as H_2_O_2_ concentration increases from 0.5 to 10 eq (20-times), it is detrimental to the enzymatic oxidation of cellulose (Fig. 4) ^15^. This is also supported by studies on the gene expressions showing that expression of LPMO is closely followed by the expression of catalase ^41^. This indicates that in nature H_2_O_2_ is controlled to be at low concentration to avoid unwanted oxidative processes inactivating the LPMOs.

Based on experimental studies, we have observed that a CuTyr-intermediate is formed when LPMO-Cu(I) and H_2_O_2_ react at room temperature. We propose that the observed CuTyr-intermediate is a real intermediate in the catalytic cycle of AA9 LPMO and therefore a key to understanding the catalytic mechanism of AA9 LPMOs which have a tyrosine in the active site (Fig. 7). The first product from this reaction between LPMO-Cu(I) and H_2_O_2_ is probably a short-lived Cu(III) intermediate as described by Margerum and co-workers ^30^. In case of no substrate, this reactive Cu(III) intermediate is transformed into the more stable CuTyr-intermediate to protect the active site from oxidation, for example in the form of alkene formation as seen in tetrapetides ^30^. In presence of substrate, it is possible that the Cu(III) intermediate can react directly with the substrate thus not forming the CuTyr-intermediate. It is yet to be established if this CuTyr-intermediate is reactive enough to abstract a hydrogen atom from a C-H bond. However, our observation on PASC oxidation supplemented with H_2_O_2_ (Fig. 4A) and of the dityrosine formation (Fig. 5) indicate that this CuTyr-intermediate acts as a powerful oxidant. We believe that the proposed CuTyr-intermediate is also important for the classical superoxo mechanism ^7,19^. In that mechanism, LPMO-Cu(I) binding to O_2_ is followed by a rapid reoxidation in an inner-sphere mechanism forming LPMO-Cu(II)-(O_2_^•─^) ^9,19,23,42^. Calculations have shown that this LPMO-Cu(II)-(O_2_^•─^) intermediate is not able to abstract a hydrogen atom from a C-H bond by itself ^7,25^. However, studies by Kjaergaard et al. (2014) have indicated that the bound superoxide radical can be replaced by water thus releasing HO_2_/O_2_^•─^ into solution which can further be transformed into H_2_O_2_ ^19^. It is known from several studies that LPMO indeed produces H_2_O_2_ in case no substrate is present ^43,44^. A possible mechanism for the reaction of reduced LPMOs with O_2_ as primary substrate (see Fig. 7 upper right) is thus a release of O_2_^−^ which disproportionates to H_2_O_2_ in solution. The formed H_2_O_2_ is then used by reduced LPMO to produce the active CuTyr-intermediate. Our stopped-flow studies indicate that LPMO-Cu(I) react very fast with H_2_O_2_.

**Fig. 7.**
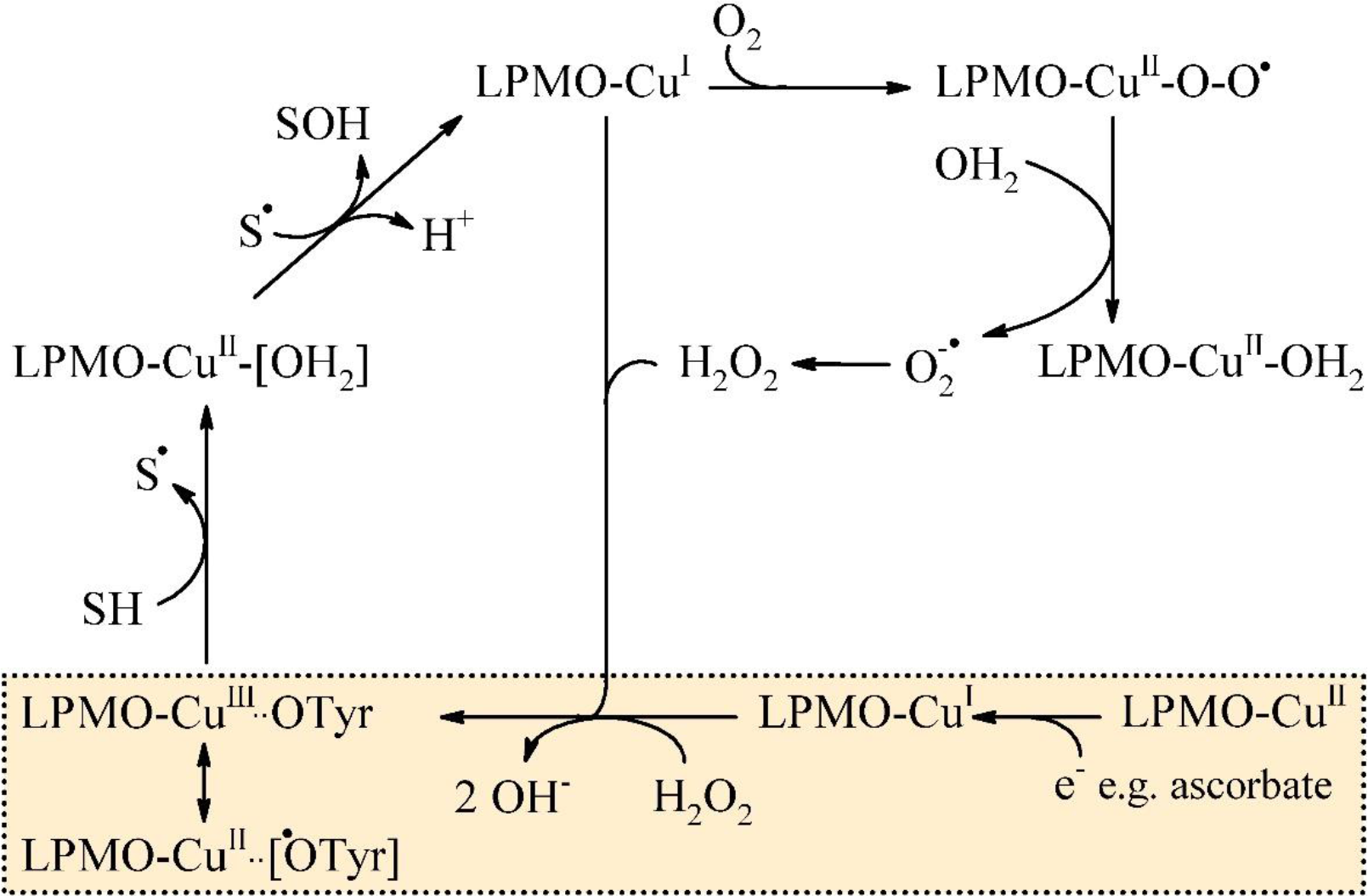
Proposed mechanism for LPMO catalyzed oxidation of polysaccharides. One electron reduction of the LPMO-Cu(II) to LPMO-Cu(I) occurs, which then reacts with H_2_O_2_ forming the CuTyr-intermediate indicated in the box. The right-side of the upper path, which is reported to produce H_2_O_2_ from the replacement of the superoxide radical ^9,19,23,26,42^ also supports the formation of CuTyr-intermediate. Then the resulting CuTyr-Intermediate could abstract a hydrogen atom from the substrate leading to hydroxylation of the substrate. The regenerated LPMO-Cu(I) can then enter a new catalytic cycle.

## CONCLUSION

In this paper, we demonstrate that Cu(I) at the active site of TaLPMO9A directly interacts with H_2_O_2_ and results in the formation of a CuTyr-intermediate. TaLPMO9A takes part in a process similar to the Cu(III) formation found by treating tri- and tetrapeptides of Cu(II) with ascorbate/H_2_O_2_ ^28,30^. However, in the case of TaLPMO9A the absorbance at 420 nm slowly disappears *without* any alkene formation but with dityrosine formation. This indicates that the active site in TaLPMO9A is protected by an additional mechanism not found in simple tri- and tetrapeptides. The reaction scheme is probably first a formation of a Cu(III) intermediate with a very short lifetime (< 1 s). This could be the active species in LPMO. In the absence of substrate, this Cu(III) species is transformed into a more stable intermediate in order to protect the active site. EPR data suggest that this long-lived intermediate could be a ferromagnetically coupled Cu(II)-^•^Tyr175 pair. However, it is still not clear if the CuTyr-intermediate is able to oxidize the substrate by itself. Our data indicate that it can react with tyrosines at the surface of TaLPMO9A resulting in the formation of dityrosine bonds. Importantly, the formation and decay of the CuTyr-intermediate do not affect the TaLPMO9A activity significantly as judged from the amplex red assay and the 2,6-DMP assay.

## METHODS

### Materials

All chemicals were of the highest purity grade commercially available and were purchased from Sigma-Aldrich unless stated otherwise. Ampliflu Red (10-acetyl-3,7-dihydroxyphenoxazine) was purchased from Cayman chemical company. H_2_O_2_ (30 %, stabilizer free) purchased from Merck Chemicals GmbH (Darmstadt, Germany). Recombinant TaLPMO9A from *T. auranticus* expressed in *Aspergillus oryzae* was donated by Novozymes A/S (Denmark).

### Purification and Copper Reconstitution

Crude TaLPMO9A from Novozymes was filtered using 0.22 μM filter and injected into a ӒKTA chromatography system equipped with Hiload 26/60 Superdex 75 prep grade column (Pharmacia Biotech) equilibrated with 20 mM MOPS buffer (pH 7.0). TaLPMO9A was eluted in single peak using MOPS buffer with the flow rate of 1 mLmin^−1^. The fractions corresponding to the single peak were pooled and concentrated using an Amicon Ultra −15 centrifugal filter (3 kDa, Merck Millipore Ltd. Ireland). Copper-(I) chloride was added to the purified TaLPMO9A in a 5mL tube on ice under anaerobic environment. After copper reconstitution, the LPMO was injected again into the ӒKTA (equipped with Hiload 26/60 Superdex 75 prep grade column) to obtain fully-loaded purified TaLPMO9A. TaLPMO9A was then concentrated and buffer exchanged to 20 mM phosphate buffer (pH 6.6) with an Amicon Ultra −15 centrifugal filter. The purity of TaLPMO9A was analyzed on sodium dodecyl sulfate-polyacrylamide gel electrophoresis (SDS-PAGE) using a Mini-PROTEAN TGX precast gel from Bio-Rad Laboratories with a gradient of 4-15%. Protein bands were visualized by staining with Coomassie blue and unstained Precision Plus Protein Standard was used for mass determination.

*apo*TaLPMO9A was prepared by dialyzing the purified TaLPMO9A extensively using D-tube Dialyzer Maxi (6-8 kDa) from Millipore (Billerica, MA, USA). Dialysis was performed in 20 mM phosphate buffer (pH 7) supplemented with 10 mM ethylenediaminetetraacetic acid (EDTA) for 12 hrs at 4 °C. Purified TaLPMO9A was loaded into a D-tube and placed in a beaker containing 600 mL ice-cold 20 mM phosphate buffer (pH 7) supplemented with 10 mM EDTA. Buffer was mixed continuously using a magnetic stirrer. During the course of dialysis, buffer was changed twice. After dialysis, the TaLPMO9A was concentrated, the buffer exchanged, and the Cu concentration was determined using inductively coupled plasma mass spectroscopy (ICP-MS).

### Copper and Protein Concentration

Protein concentration of purified TaLPMO9A was determined by its absorbance at 280 nm using a NanoDrop spectrophotometer from Fisher Scientific. Extinction coefficient of TaLPMO9A was calculated from TaLPMO9A sequence using online tool available at https://web.expasy.org/protparam/. Copper concentrations were determined using high resolution inductively coupled plasma mass spectroscopy (ICP-MS) on an Aurora M90 ICP-MS system from Bruker. For this, TaLPMO9A was diluted in 1.5% nitric acid. Each sample was measured in triplicate and average values are reported.

### LPMO activity assay

Horseradish peroxidase (HRP)-coupled Amplex red assay was performed on SpectraMax M2 multi-detection microplate reader using black 96-well plate with clear bottom from Nunc ^44^. All the reactions were performed in 20 mM sodium phosphate buffer, pH 6.0 at 37 °C. 200 μL of reaction mixture contained 100 μM EDTA, 80 μM ascorbate, 50 μM ampliflu red, 20 UmL^− 1^ HRP and, 2.0 μgmL^−1^ LPMO. Reaction was initiated by adding TaLPMO9A and resorufin fluorescence was recorded with an excitation and emission wavelength of 557 nm and 583 nm, respectively. Fluorescence was measured for 10 min with 5 sec shaking before the measurement. A linear relation between fluorescence and H_2_O_2_ concentrations in the range of 0.1 – 2 μM H_2_O_2_ was observed and the slope (28450 counts μmol^−1^) was used for the calculation of an enzyme factor to convert the fluorimeters readout (counts min^−1^), into enzyme activity. LPMO activity was defined as one μmol H_2_O_2_ generated per minute under the defined assay conditions.

LPMO activity assay using 2,6-dimethoxyphenol (2,6-DMP) was performed as described by Breslmayr et al. (2018) ^34^. The standard reaction mixture for the LPMO activity assay contained 100 μM H_2_O_2_, 1.0 mM 2,6-DMP and 4 μM of TaLPMO9A. Reaction was performed in phosphate buffer (pH 7.5) at room temperature. Reaction was initiated by adding TaLPMO9A and UV-vis spectra was recorded every 30 seconds for 10 min on Agilent UV– visible ChemStation.

### LPMO oxidation

LPMO oxidation was achieved by mixing the LPMO with equimolar amounts of L-ascorbate and various concentrations of H_2_O_2_ ^30^. LPMO was prepared in 20 mM phosphate buffer (pH 6.6) while ascorbate and H_2_O_2_ stock solutions were prepared in deionized filtered (0.22 μm) Milli-Q water. The concentration of H_2_O_2_ was confirmed by UV spectroscopy based on the absorption at 265 nm (ε 10 M^−1^cm^−1^) ^45^. UV-Vis spectral change was recorded using a SHIMADZU UV-3600 UV-VIS-NIR spectrophotometer. Spectra were recorded from 300 to 700 nm wavelength. All the solutions were freshly prepared and stored in dark tubes on ice prior to the experiment.

### Detection of dityrosine

TaLPMO9A was oxidized by mixing equimolar concentrations of L-ascorbate and H_2_O_2_. Change in the intrinsic fluorescence of TaLPMO9A due to the formation of dityrosine was recorded at 340-500 nm after excitation at 320 nm ^33^, on a Jasco FP-6300 spectrofluoromater using a 1 cm cuvette. Spectral changes were monitored at different time points for 24 hours.

### EPR Spectroscopy

EPR spectra were recorded for reduced, oxidized and unmodified *apo*TaLPMO9A and *holo*TaLPMO9A on a Bruker Elexsys E500 equipped with a Bruker ER 4116 DM dual mode cavity spectrometer equipped with a SuperX CW-EPR bridge, and a nitrogen flow cryostat (Helitran; Advance Research Systems). EPR spectra were recorded at room temperature as well as 77 K using microwave frequency, 9.63 GHz; microwave power, 10 milliwatts; modulation amplitude, 8 G; and attenuation 10 dB. For room temperature measurements, samples were loaded in capillary tube and then placed into EPR quartz tubes. Theoretical EPR spectra were fitted to the experimental ones as detailed in ref ^46^. To reproduce the Cu^2+^ signals we used the spin Hamiltonian

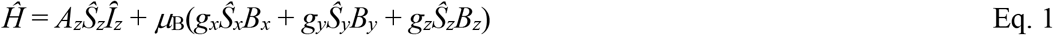

with S = 1/2 and I = 3/2 pertinent to the electronic spin doublet and the Cu nucleus, respectively. Initially, we included in the spin Hamiltonian the *x* and *y* components of the hyperfine interaction. However, the experimental data did not warrant such a detailed analysis. Therefore, equation (1) was used in the fitting of all Cu^2+^ signals with axial *g* factors, i.e. with g_x_ = g_y_.

### LPMO catalyzed oxidation of PASC

PASC microcrystalline cellulose substrate was prepared by treating avicel with phosphoric acid as described by Wood et al. (1988) ^47^. 4 grams of Avicel was dispersed into 100 mL of phosphoric acid (86 % w/v) at 60 °C and magnetically stirred for an hour. 1900 mL Milli-Q water was slowly dripped into the solution under stirring condition. Then suspension was transferred to 4 °C for sedimentation. After sedimentation, the supernatant was removed and the suspension was washed four times with Milli-Q water. The solution was neutralized using Na_2_CO_3_.

TaLPMO9A catalyzed oxidation of PASC contains TaLPMO9A (0.1 mM), ascorbate (2 mM), and PASC (0.4 %, w/v)^48^. Reaction was carried out in absence or presence of H_2_O_2_ (0.1 mM / 2 mM) in 20 mM phosphate buffer (pH 6.0) at 25 °C for 30 min. Reaction was stopped using 0.1 M NaOH and then samples were centrifuged. Supernatant was separated, filtered (0.22 μM) and analyzed for oligosaccharide by HPAEC. HPAEC was run on an ICS5000 system equipped with a PAD detector (Dionex, Sunnyvale, CA, USA) with a CarboPac PA1 column (2 × 50 mm guard column followed by a 2 × 250 mm analytical column).

### High-Performance Anion Exchange Chromatography (HPAEC)

HPAEC analysis of the released oligosaccharides from the cellulosic substrate was performed on an ICS-5000+ system, equipped with a PAD detector (Thermo Scientific), with a CarboPac PA1 column (two 2 × 50 mm guard columns followed by a 2 × 250 mm analytical column), and operated at 0.25 ml min^−1^ and 30°C. The chromatographic separation of aldonic acids was carried out as described by Westering et al. (1913) ^49^. For the elution the following gradient was applied (Eluent A, 0.1M NaOH; Eluent B, 1 M NaOAc in 0.1 M NaOH): 100% A:0% B to 90% A:10% B (10 min), then to 70% A:30% B (25 min) and lastly 0% A:100% B (30 min). For reconditioning of the column 100% A:0% B was applied for 15 min (39-54 min). The cello-oligosaccharide peaks were annotated according to elution profiles of commercially available non-oxidized oligosaccharides (DP2-DP6). The C4-oxidized glucose (C4-keto aldose) peaks appeared after 25 min.

### LPMO Stopped-flow kinetics

Fast kinetic studies were performed using a π*−180 stopped-flow spectrometer (Applied Photophysics, Leatherhead, UK) with an observation cell of the light path length of 2mm.

Calibrations of the zero time-point and dead-time (the time from the zero time to the first good data point) were done by mixing 2,6-Dichlorophenolindophenol (DCIP) with different concentrations of excess L-ascorbic acid (AA) (under pseudo-first order reaction conditions) at a 1:1 ratio (v:v). Absorption was measured at 524 nm, data acquisition and analysis were done using the Pro-Data software (Applied Photophysics). The zero time-point and dead-time may change with AA concentration, and all analysis were based on kinetic traces that were adjusted to actual time points of mixing determined by DCIP–AA experiments at different AA concentrations. The experimental data were fitted to single exponential equations. All the initial data points were omitted from the traces until satisfactory fits to single exponentials were obtained, and a common point of intersection was achieved for a range of AA concentrations. The point of intersection of the fitted curves is the true zero time for the mixing.

Kinetic studies of fast reactions between LPMO, AA and H_2_O_2_ were performed and the reaction species were followed at 420 nm with time scales from 60 to 1000s at 20 °C. The first stopped-flow syringe contained reduced TaLPMO9A, and second syringe contained varying concentrations of H_2_O_2_ (0.05 – 3.0 mM). The fit to the data was made between 5-1000 s thereby removing the rising part of the transients from the fit. The fit was made using the standard double exponential equation (Eq. 2).

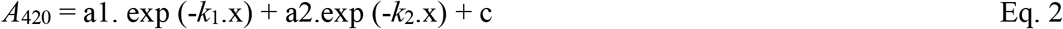

where *k*_1_, and *k*_2_ are the observed rate constants for the fast and slow phases respectively, a1 and a2 are the relative amplitude values for the two phases, and c is an offset value to account for a non-zero baseline. Baseline tend to fluctuate during multiple shots. Therefore, baselines were normalized. All experiments were done in duplicates for each reactant concentration. After rapid mixing, absorbance changes were monitored at a 420 nm and appropriate controls were recorded in all cases to exclude the possibility of artifacts.

## Acknowledgments

This work is supported by the Novo Nordisk Foundation project “Harnessing the Energy of the Sun for Biomass Conversion” (NNF16OC0021832). We also acknowledge the donation of enzymes from Novozymes A/S. All the data relevant to this study are included in the main paper or the supplementary materials.

## Authors contributions

M. J. B. and R. K. S. designed, performed and analyzed most of the experiments. R. S. performed the stopped-flow kinetics experiments. B. M. and H. W. helped to perform and analyze the PASC oxidation and EPR spectroscopy data, respectively. B.v.O helped to analyse the stopped-flow data. M. J. B. and R. K. S. wrote the manuscript with support from C. F. and D. A. R. All authors participated in critical analysis of the data and finalizing the manuscript.

## Competing financial interests

The authors declare no competing financial interests.

