## Supplementary Information for "Detection and characterization of a novel copper-dependent intermediate in a lytic polysaccharide monooxygenase"

### FOR

**Table S1. Stopped-flow kinetic data.** Kinetic parameters fitted to the raw data as a function of varying $H_2O_2$ concentration for reduced TaLPMO9A. Data was analyzed using the Pro-Data software (Applied Photophysics).

| **TaLPMO9A (mM)** | **Ascorbate (mM)** | **$H_2O_2$ (mM)** | **$[H_2O_2]$/[TaLPMO9A]** | **$k_1$ ($s^{-1}$)** | **$k_2$ ($s^{-1}$)** | **a1** | **a2** | **c** |
|---|---|---|---|---|---|---|---|---|
| 0.203 | 1 | 0.25 | 1.23 | 0.00280 | 0.00140 | 0.00167 | 0.00129 | 0.00562 |
| 0.203 | 1 | 0.5 | 2.46 | 0.00797 | 0.00291 | 0.00263 | 0.00323 | 0.00400 |
| 0.203 | 1 | 1.0 | 4.93 | 0.01104 | 0.00256 | 0.00323 | 0.00392 | 0.00294 |
| 0.203 | 1 | 1.5 | 7.39 | 0.01313 | 0.00260 | 0.00571 | 0.00245 | 0.00337 |
| 0.203 | 1 | 2.0 | 9.85 | 0.01715 | 0.00235 | 0.00567 | 0.00410 | 0.00120 |
| 0.203 | 1 | 2.5 | 12.3 | 0.01791 | 0.00116 | 0.00700 | 0.00421 | 0.00170 |
| 0.203 | 1 | 3.0 | 14.8 | 0.01966 | 0.00092 | 0.00747 | 0.00731 | 0.00062 |

**Table S2. EPR data for experiments performed at 77K.** I, II and III denote TaLPMO9A, reduced TaLPMO9A, and reduced TaLPMO9A + $H_2O_2$, respectively. Substrate represents the addition of 0.1% PASC to the reaction mixture. Numbers in brackets represent the respective standard deviation.

| **Parameter** | **No substrate** | | | **Substrate** | | |
|---|---|---|---|---|---|---|
| | **I** | **II** | **III** | **I** | **II** | **III** |
| $A_z$ (MHz) | 492(1) | 493(2) | 495(2) | 493(1) | 492(1) | 493(2) |
| $g_z$ | 2.2718(3) | 2.2725(4) | 2.2730(5) | 2.2722(3) | 2.2722(3) | 2.2718(4) |
| $g_{xy}$ | 2.0558(1) | 2.0563(2) | 2.0569(2) | 2.0561(1) | 2.0564(1) | 2.0563(2) |
| Cu (conc.) | 1792(45) | 1504(54) | 1381(56) | 999(31) | 802(25) | 773(27) |

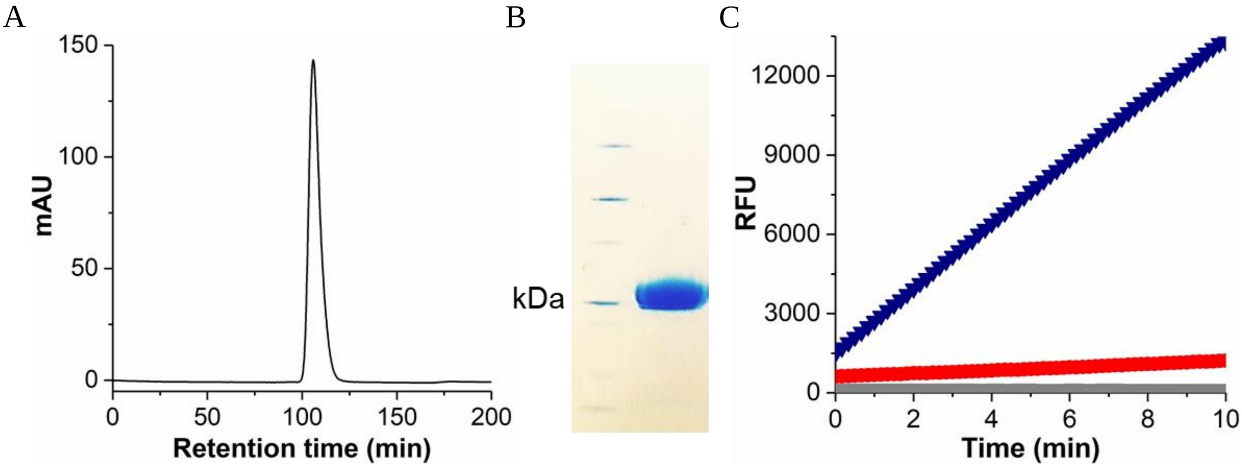


**Fig. S1. Purification and activity of TaLPMO9A.** (**A**) Chromatogram showing the last step of Cu (I) reconstituted TaLPMO9A purification using size exclusion chromatography. (**B**) SDS-PAGE analysis of purified TaLPMO9A. (**C**) Plot showing activity measurement of purified TaLPMO9A (▼) using amplex red assay. Red solid circle (●) and gray solid square indicate the reaction mixture without TaLPMO9A and buffer, respectively. Reaction was carried out in 20 mM phosphate buffer containg ascorbate (0.08 mM), EDTA (0.1 mM), Amplex red (0.05 mM), HRP (20 U), TaLPMO9A at 37 ◦C.

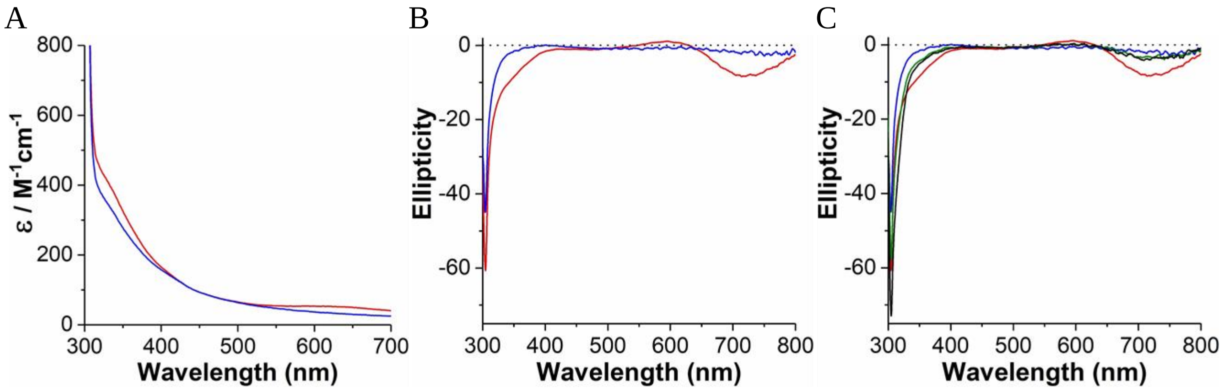


**Fig. S2. Absorption spectrum of *holo*TaLPMO9A before and after addition of ascorbate.** (**A**) Vis-spectra of TaLPMO9A (1.93 mM) corrected for baseline. Copper(II) signal at ~340 nm (shoulder) and ~640 nm (peak). (**B**) The CD spectrum of the *holo*TaLPMO9A has a positive band at 600 nm and two broad negative bands at 305 and 720 nm. Upon adding ascorbate, the bands at 600 and 740 nm disappear, while 305 nm band appears to lose the shoulder band at 350 nm. (**C**) CD spectra of *holo*TaLPMO9A upon addition of ascorbate and $H_2O_2$. Red line and blue line represent *holo*TaLPMO9A and reduced *holo*TaLPMO9A, respectively. Green and black lines represent reduced TaLPMO9A + $H_2O_2$ at 60 s and 5 min, respectively. CD spectra were recorded at room temperature using a 10 mm cuvette.

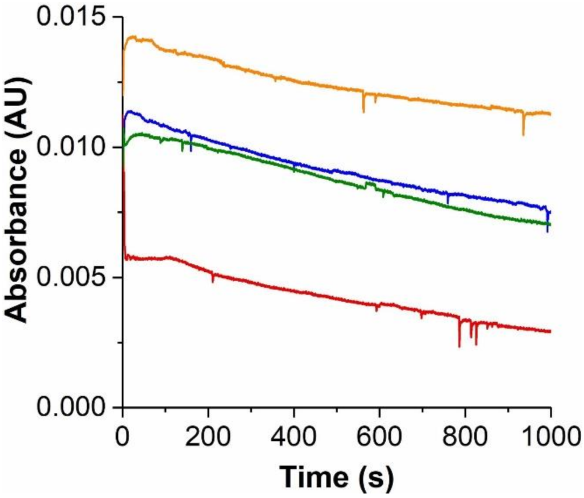


**Fig. S3. Single wavelength absorption transients at 420 nm for the reaction of reduced TaLPMO9A with $H_2O_2$ at different pH.** (**B**) Transients for change in absorption at 420 nm observed for 1000 s at pH 5.9, 6.3, 7.1 and 7.6. Red, blue, orange and green lines represent the absorbance at pH 5.9, 6.3, 7.1 and 7.6, respectively. Experimental conditions: *holo*TaLPMO9A 0.2 mM, ascorbate 1 mM, and 0.4 mM $H_2O_2$ in 20 mM phosphate buffer, 20 $^oC$.

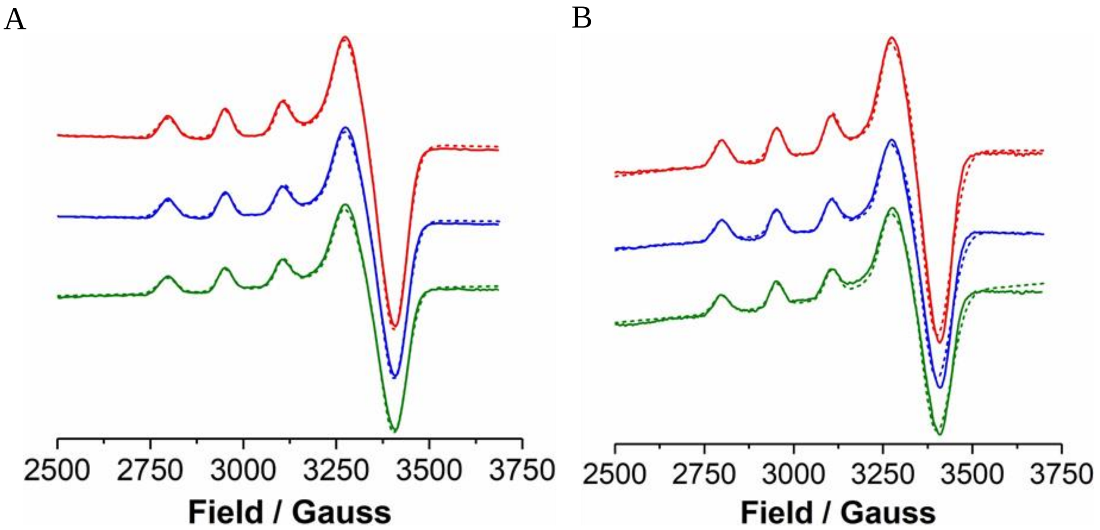


**Fig. S4. EPR spectra of *holo*TaLPMO9A.** (**A**) X-band EPR spectra of TaLPMO9A (red line), reduced TaLPMO9A (blue line) and $H_2O_2$ treated reduced TaLPMO9A (green line) at 77 K in 20 mM phosphate buffer, pH 6.6. Experimental conditions: TaLPMO9A 1.7 mM, ascorbate 2 mM, $H_2O_2$ 1 mM at pH 6.6 in 20 mM phosphate buffer, at room temperature. (**B**) X-band EPR spectra of TaLPMO9A (red line), reduced TaLPMO9A (blue line) and $H_2O_2$ treated reduced TaLPMO9A (green line) in presence of substrate (PASC) at 77 K in 20 mM phosphate buffer, pH 6.6. All the spectra were corrected for background, which was recorded with pure buffer solution in EPR tube. Experimental and calculated spectra are indicated with solid and dashed lines, respectively.

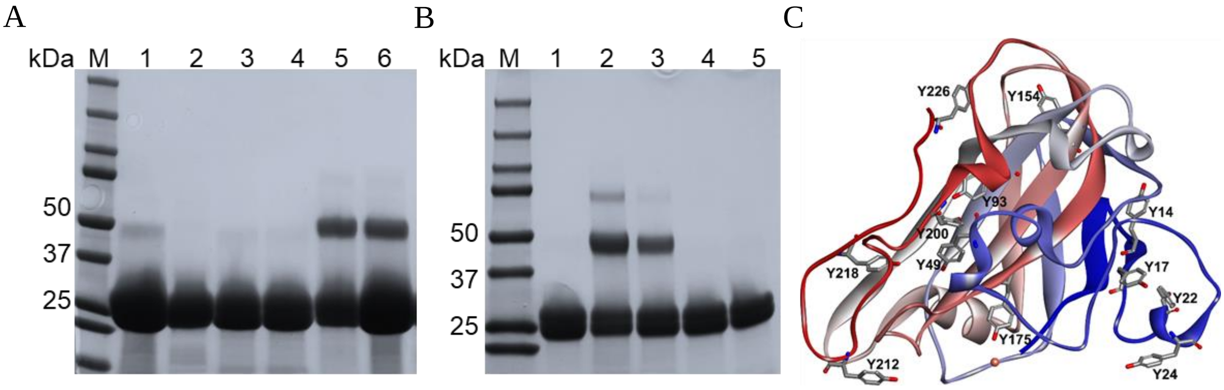


**Fig. S5. The appearance of TaLPMO9A dimers correlates with the formation of dityrosine.** (**A**) SDS-PAGE analysis of the TaLPMO9A after initial intermediate formation. Lane 1: Purified *holo*TaLPMO9A (0.4 mM), Lane 2: *holo*TaLPMO9A (0.2 mM) + 1 mM ascorbate, Lane 3: *apo*TaLPMO9A (0.2 mM) + 1 mM ascorbate, Lane 4: *apo*TaLPMO9A (0.2 mM) + 1 mM ascorbate + $H_2O_2$, Lane 5: *holo*TaLPMO9A (0.2 mM) + 1 mM ascorbate + $H_2O_2$, Lane 6: *holo*TaLPMO9A (0.4 mM) + 1 mM ascorbate + $H_2O_2$. (**B**) Lane 1: *holo*TaLPMO9A + $H_2O_2$, Lane 2: *holo*TaLPMO9A + 1 mM ascorbate + $H_2O_2$, Lane 3: *holo*TaLPMO9A + 1 mM ascorbate + $H_2O_2$, Lane 4: *holo*TaLPMO9A (0.2 mM) + 1 mM ascorbate, Lane 5: *holo*TaLPMO9A. (**C**) Crystal structure of TaLPMO9A (PDB: 2YET[5]). Tyr residues and Cu (II) are shown with stick and ball model, respectively.

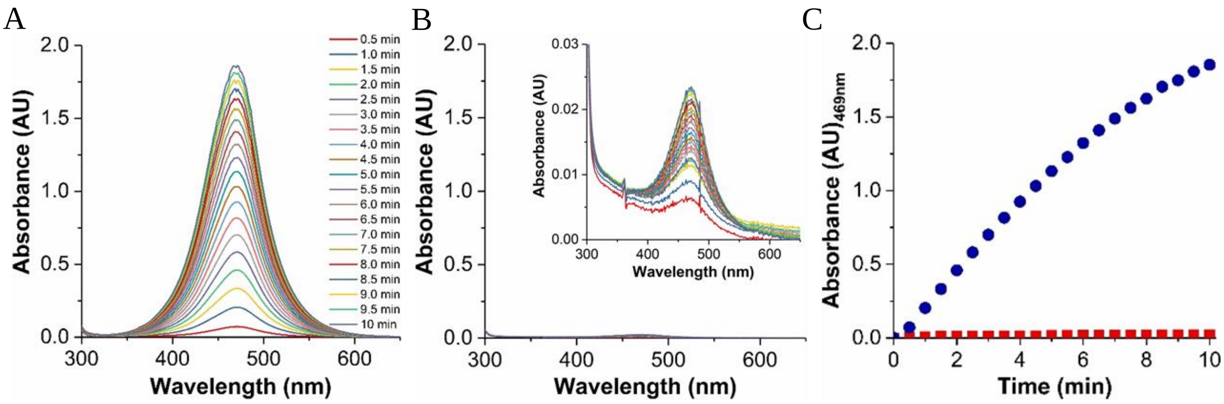


**Fig. S6. 2,6-dimethoxyphenol assay for TaLPMO9A.** (**A**) LPMO activity assay was performed at room temperature in phosphate buffer (100 mM, pH 7.5). $H_2O_2$ concentration of 100 μM, a 2,6-DMP concentration of 1.0 mM and TaLPMO9A (4 μM). (**B**) LPMO activity assay was performed at room temperature in phosphate buffer (100 mM, pH 7.5). 2,6-DMP concentration of 1.0 mM and TaLPMO9A (4 μM). Inset shows enhanced spectra of TaLPMO9A activity in absence of $H_2O_2$. (**C**) Comparison of absorbance at 469 nm for TaLPMO9A catalyzed activity in presence (blue solid circle) and absence of $H_2O_2$ (red solid square) over time.

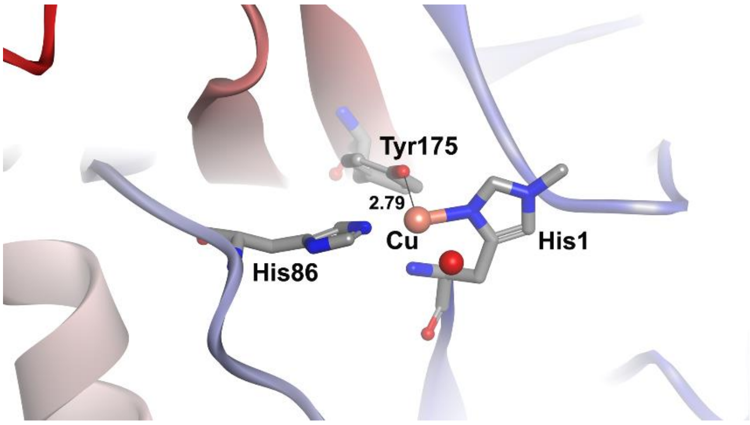


**Fig. S7. The crystal structure of TaLPMO9A showing the active site environment.** Tyrosine (Try175) and histidine residues are indicated with stick model. Cu (II) is shown with an orange sphere (PDB: 2YET).

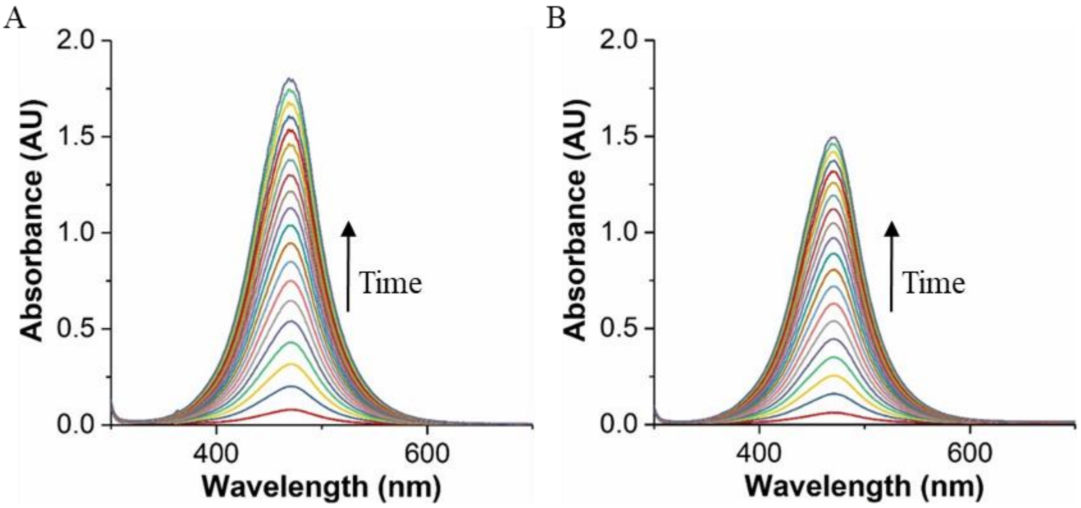


**Fig. S8. 2,6-dimethoxyphenol assay for oxidized TaLPMO9A.** LPMO activity assay for the oxidized TaLPMO9A. TaLPMO9A oxidation was done using 0.5 mM $H_2O_2$/ 1mM ascorbate (**A**) and 10 mM $H_2O_2$/ 1mM ascorbate (**B**). LPMO activity was then measured using the 2,6-dimethoxyphenol assay. The assay was performed for 10 minutes by recording spectra every 30 seconds at room temperature in phosphate buffer (100 mM, pH 7.5). 2,6-DMP concentration of 1.0 mM and TaLPMO9A (4 μM). Oxidized TaLPMO9A was purified and buffer was changed to remove residual ascorbate and $H_2O_2$ before assay for activity. The activity in A is calculated to be 91 % of *holo*TaLPMO9A activity while activity in B is 78 %.
